# Mapping SARS-CoV-2 Nucleocapsid Function with Nanobodies

**DOI:** 10.64898/2026.01.26.701894

**Authors:** Jules B. Reyes-Weinstein, Timothy A. Bates, Aidan Anastas, Ayna Alfahdli, Mila Trank-Green, Savannah K. McBride, Michelle Garcia, Matthew A. H. Parson, Meredith L. Jenkins, John E. Burke, Eric Barklis, Fikadu G. Tafesse

**Author notes:** Correspondence: Fikadu G. Tafesse. These authors contributed equally (Jules B. Reyes-Weinstein, Timothy A. Bates).

## Abstract

The SARS-CoV-2 nucleocapsid (N) protein is essential for viral RNA packaging, replication, and immune modulation. Despite its central role, the mechanistic contributions of its individual domains, the N-terminal domain (NTD), C-terminal domain (CTD), and the intrinsically flexible linker (LINK), remain poorly defined, largely due to the protein’s structural complexity. In this study, we developed a panel of twelve alpaca-derived nanobodies (VHHs) targeting the NTD, CTD, and LINK regions of N. Using ELISA and biolayer interferometry, we characterized their binding affinities, and we mapped their epitopes via hydrogen-deuterium exchange-mass spectrometry (HDX-MS) and structural modeling. When expressed intracellularly, these VHHs inhibited SARS-CoV-2 infection. In vitro, they disrupted phase separation of the N protein, a critical step in viral replication. Strikingly, VHHs targeting each domain independently blocked both phase condensation and viral replication, underscoring the functional importance of all three regions. These findings establish domain-specific VHHs as versatile tools for dissecting N biology, with promising therapeutic potential.

**Importance:** SARS-CoV-2 and emerging coronaviruses remain a major global health threat, yet critical gaps persist in our understanding of their molecular pathogenesis. The nucleocapsid (N) protein, the most abundantly expressed SARS-CoV-2 antigen, plays essential roles beyond genome packaging, including immune evasion and intracellular organization. Here, we generate and characterize a panel of domain-specific nanobodies (VHHs) that enable precise dissection of N’s functional architecture. Using integrated biochemical, structural, and virological approaches, we uncover distinct mechanisms of viral inhibition, including disruption of phase condensation through a conserved linker region. These findings address long-standing knowledge gaps about a multifunctional viral protein and establish VHHs as powerful, modular tools for probing coronavirus biology, with broad potential for therapeutic, diagnostic, and cell biology applications.

## Introduction

The COVID-19 pandemic dealt a devastating blow to global health and economic systems, and SARS-CoV-2 has now become a permanent part of human society. The pandemic potential of coronaviruses have been appreciated since the mid 1990’s, and it is likely that future zoonotic spillover events will involve similar coronaviruses^1,2^. Decades of previous work on the basic biology and pathogenic mechanisms of coronaviruses helped speed the development of the life-saving vaccines and therapeutics that blunted the worst impacts of the pandemic, but in order to create more broadly useful anti-coronaviral preventatives and treatments we need to better understand the many distinct functional roles of SARS-CoV-2 structural proteins, such as Nucleocapsid (N).

As with other CoVs, the 30 kb positive-sense RNA genome of SARS-CoV-2 encodes four structural proteins: Spike, Nucleocapsid, Membrane, and Envelope^3,4^, each of which plays an essential role in viral replication. While viral entry is generally mediated by the interaction of S with the human ACE2 receptor and cleavage by TMPRSS2 protease^5^, virions can also enter through direct fusion of the plasma membrane if S is sufficiently primed^6^. Post membrane fusion, the N-coated viral genome is released into the cytoplasm where it sheds the N protein and traffics to the endoplasmic reticulum (ER) by poorly understood mechanisms, associates with the host ribosomal ribonuclear complex, and is subsequently translated^7–9^. The SARS-CoV-2 genome encodes several structural and non-structural proteins (NSP), the expression of which facilitates the formation of double membrane-bound replication organelles^10^, called replication translation complexes (RTC), by manipulating the host ER membrane. These sites serve numerous purposes: sequestering viral material from the host, keeping viral NS proteins and RNA in close association, and concentrating structural proteins within the membrane so that they can go on to form the viral envelope. The N protein can both multimerize and complex with newly synthesized RNA genomes which together drive liquid-liquid phase separation (LLPS), forming densified regions that further separate the viral genomes from the rest of the cytosolic milieu^11,12^. These phase-separate globules are then loaded into nascent viral particles which, after further maturation, are then secreted through the Golgi and ultimately released through extra-cellular vesicles.

The N protein contains several ordered and disordered domains (**Figure 1A**). The ordered N-terminal domain (NTD) and C-terminal domain (CTD) are flanked by three disordered domains, the N-terminal unstructured N-arm domain, the C-terminal unstructured C-tail domain, and a flexible linker (LINK) segment between NTD and CTD^7,12^. The N protein natively forms dimers when purified; and is believed to form higher-order oligomers, particularly in the presence of RNA^7,12^. Due to the numerous disordered regions, domains have only been crystalized individually, with the NTD crystalized as a monomer and the CTD in its native dimer form^13,14^. In previous studies of N from SARS-CoV-1, the NTD was primarily implicated in RNA-binding and the CTD in dimerization^15,16^. However, further research has revealed more redundant functionality between the NTD and CTD, while LINK also appears to play a structural role despite its lack of intrinsic structure. It has been shown that N LINK residues 210-246 were conserved across multiple CoV species and were essential for dimerization^17^. Mutagenesis experiments using SARS-CoV-2 virus-like particles (VLP) indicate that mutations in LINK are more likely to increase packaging efficiencies than those in NTD or CTD^18^. The N protein also has secondary functions outside of its structural and replicative roles including modulation of host immune genes and stress responses^19–22^. These secondary functions, which drive pathogenesis and enhance replication, are believed to be due to non-specific nucleic acid binding.

**Figure 1:**
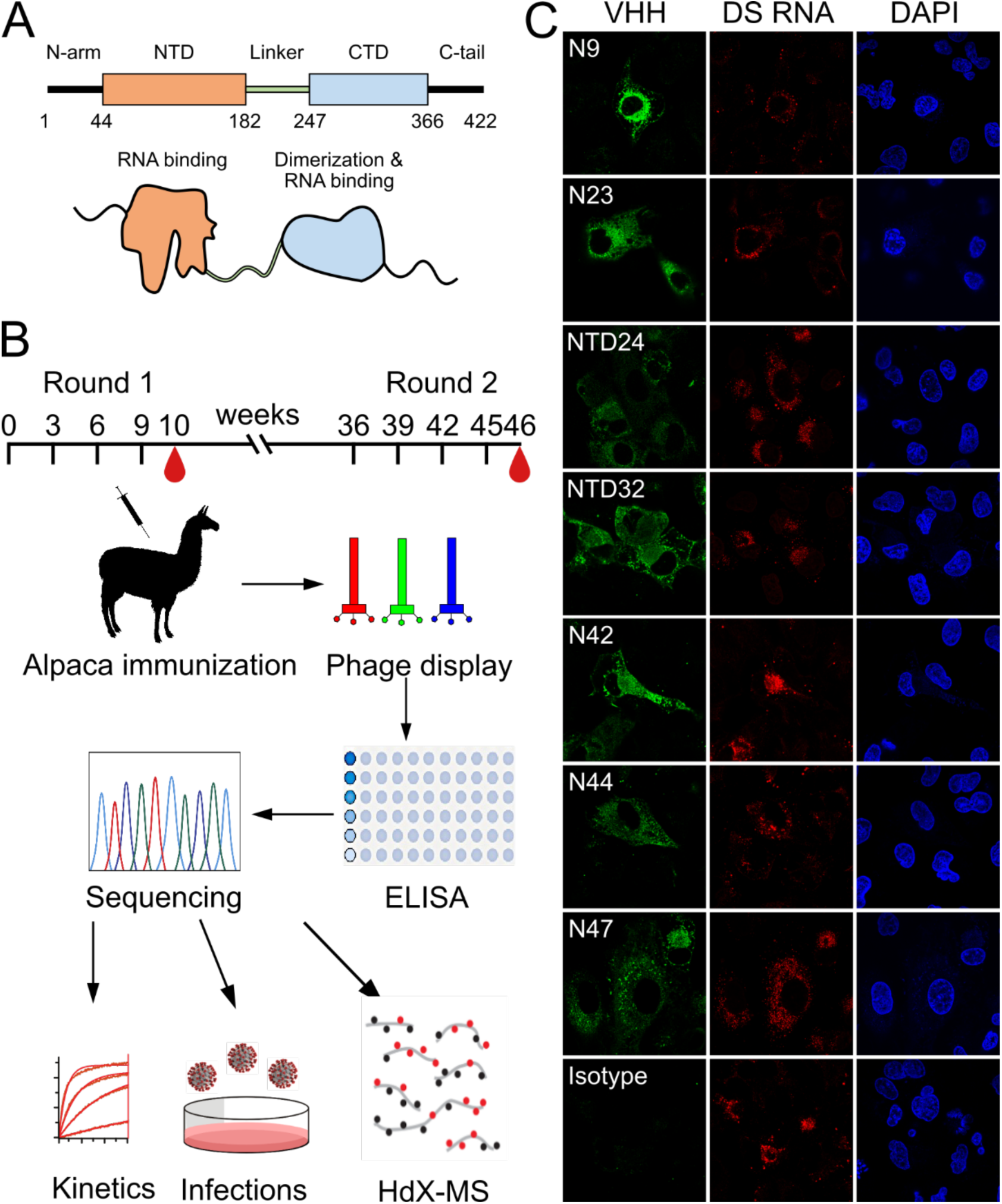
Phage display discovery of anti-nucleocapsid nanobodies. A) Schematic of the nucleocapsid (N) protein. The disordered N-arm and C-tail flank the structured N-terminal domain (NTD) and C-terminal domain (CTD) which are attached via a flexible linker (LINK) region. The NTD and CTD can both bind RNA while only the CTD can form homodimers with other N proteins. B) Overview of alpaca immunization schedule and workflow. Alpacas were immunized with two series of recombinant SARS-CoV-2 N injections. VHH mRNA was harvested from PBMCs, amplified by PCR, expressed in bacteriophage, and selected for binding strength by phage display. Enriched sequences used in downstream applications. C) Immunofluorescence of 2 µg/ml solutions of each high-binding VHHs in A549-ACE2 cells infected with WA1 at an MOI of 0.1. Cells stained for double-stranded (DS) RNA (red), nanobody (green), and DAPI (blue). Isotype control is specific for influenza.

Despite progress in characterizing the basic functions of N across CoVs, many mechanistic details remain unresolved. Structural analysis is hampered by the protein’s high degree of intrinsic disorder, preventing full-length crystallization or cryo-EM studies. While whole viral particles have been imaged by cryo-EM/ET, the packaging of the viral genome has so far proven too variable to allow generation of suitable class averages for solving residue-level structures. NMR studies have given some information about which residues are involves in nucleic acid binding^23^, and truncation variant studies have implicated various portions of the N protein in assembly of ribonucleosomes^24^, however full length structures of N, or N-RNA complexes remain unsolved, and the functional redundancy of its domains makes interpretation of experimental results less straightforward.

To address these gaps, we developed a panel of high-affinity alpaca-derived nanobodies, also called variable heavy chain antibody fragments (VHHs), targeting the NTD, CTD, and LINK regions of the SARS-CoV-2 N protein. Unlike conventional antibodies, VHHs can fold and function in the reducing environment of the cytoplasm, making them ideal for probing intracellular protein dynamics and phase transitions^25,26^. This feature is particularly advantageous for targeting SARS-CoV-2 N, which is both structurally dynamic and sensitive to redox conditions.

Here, we present 12 VHHs that map to distinct N protein domains. We use ELISA and BLI to characterize their binding affinities; we use hydrogen-deuterium exchange mass spectrometry (HDX-MS) and *in silico* structural modeling to assigned their epitopes; we use *in vitro* phase condensation assays to demonstrate domain-specific functional disruption; and we perform inhibition assays with real SARS-CoV-2 to evaluate which VHHs can inhibit viral replication. We observe an interesting relationship between the ability to disrupt phase condensation and the ability to block viral replication, and we found that LINK-specific VHHs had an outsized impact on both LLPS and viral replication. These findings establish domain-specific VHHs as versatile tools for dissecting N biology and underscore the complexity and interdependence of N’s functions. Our study also opens new avenues for designing antiviral therapeutics that prevent replication by interrupting ribonuclesome assembly.

## Results

### Generation of a diverse panel of N-specific VHHs

To generate VHHs targeting the SARS-CoV-2 N protein, we immunized an alpaca with recombinant full-length N for two rounds of four doses (**Figure 1B**). Following immunization, a VHH library was constructed from peripheral blood lymphocytes and subjected to phage display panning against the full-length N protein. We identified six enriched VHH sequences from the initial round of immunization, and an additional six higher-affinity VHHs after the second round (**Supplemental Table 1**).

To evaluate *in situ* binding, we infected A549-ACE2 cells with the original WA1/2020 SARS-CoV-2 strain (WA1) and stained them with the top seven VHH candidates (**Figure 1C**). All candidates successfully stained infected cells, in contrast to our influenza-specific isotype control VHH.

To properly understand the binding kinetics of our VHH candidates we employed two approaches to quantify binding strength: ELISA to measure their 50% maximal binding concentrations (EC_50_), and BLI to measure their dissociation constants (*K_D_*). Based on ELISA data, the second immunization round produced substantially higher affinity VHH candidates than the first round (**Supplemental Table 1, Supplemental Figures 1-2**). The highest affinity first round candidate, VHH N23, had an ELISA EC_50_ of 273.9 nM against the full-length N used in immunization. In contrast, the highest affinity second round candidate, NTD32, had an EC_50_ of 60 pM, over a 4000-fold improvement (**Figure 2A**). These findings were supported by our BLI results (**Figure 2B-C**). N23 did not bind strongly enough to yield a *K_D_* in BLI, whereas NTD24 had a *K_D_* of 63 pM. It is well-established that avidity can improve overall binding strength by several orders of magnitude^27^, which we tested by constructing a human IgG1-based Fc-conjugated bivalent VHH for each of our low-affinity VHHs^28^. The Fc-conjugated variant of VHH23 had an EC_50_ of 0.3 nM, about a 1000-fold improvement. Furthermore, it could now bind effectively in BLI, yielding a *K_D_* of 2.78 nM (**Supplemental Figures 1-2**). However, the second round of immunization generated VHH clones with sub-nanomolar affinities as monomers, obviating the need for Fc-multimerization, which requires mammalian expression. Because of this, we focused our attention on those VHHs with high affinity as monomers.

**Figure 2:**
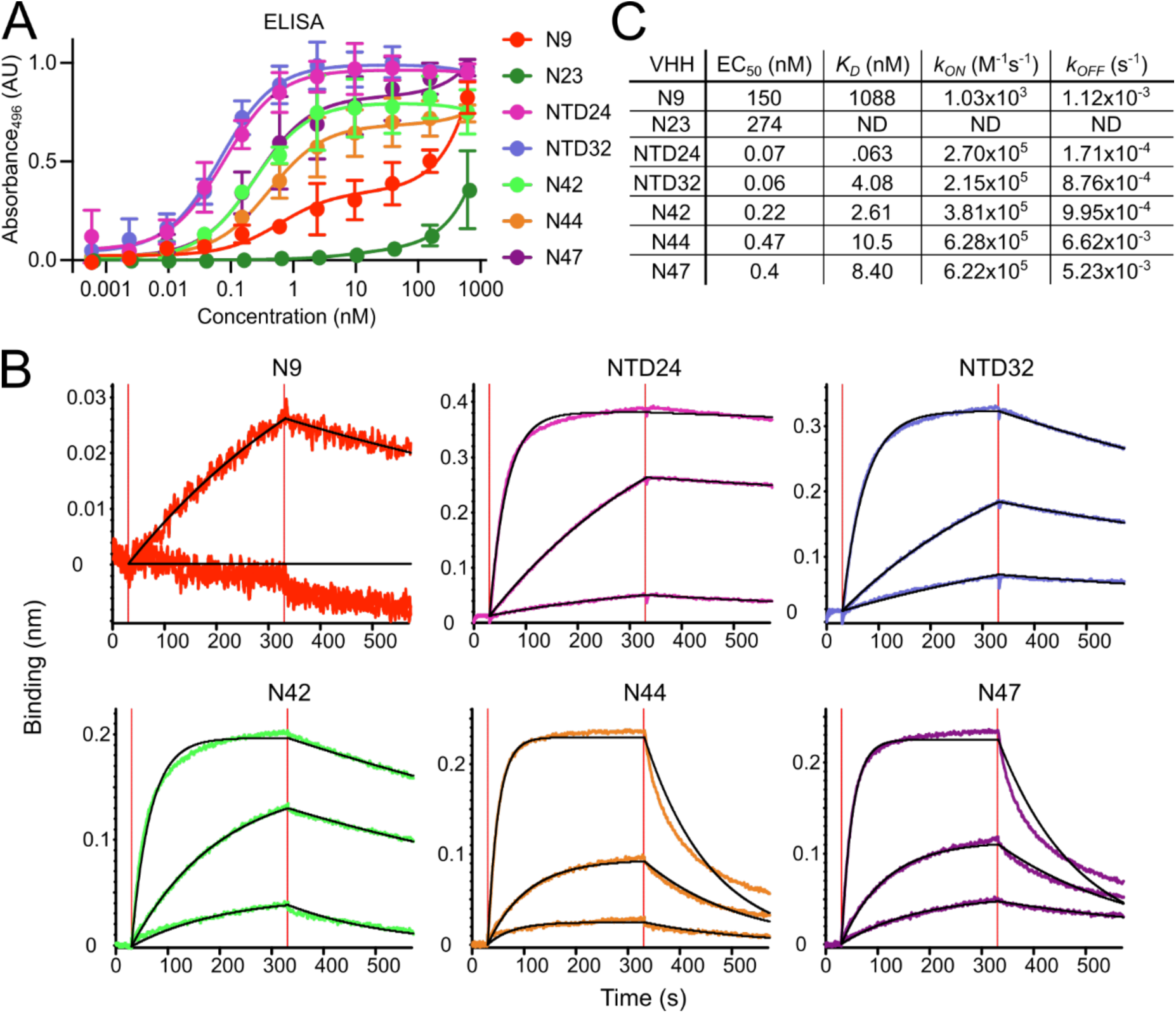
Additional alpaca immunization raised VHHs of high affinity against full-length. **N.** A) ELISA plots showing each VHH at various concentrations bound to 1 µg/ml SARS-CoV-2 N and stained with anti-nanobody antibody (3 experimental replicates). VHH N23 was from the first immunization while N9, NTD24, NTD32, N42, N44, and N47 were from the second. B). Biolayer-interferometry schematic and sample readout showing streptavidin tips bound with 1 µg/ml of biotinylated N, then 100nM of VHH (association). Dissociation was measured by removing VHHC) Summary data of EC_50_ calculated from ELISA data, *K_D_*, *k_ON_*, *k_OFF_* calculated from BLI. “ND” for not determined.

### Determination of binding epitopes of N-specific VHHs

We next sought to identify where these VHHs bound on the N protein. To accomplish this, we first carried out a competitive BLI method to determine which clones bound overlapping epitopes (**Figure 3A**), using our previously published method^29^. We first bound one VHH before incubating with a second VHH, without a wash step. A rising signal from the second VHH indicates a distinct non-overlapping epitope, while a flat curve implies a shared epitope or a rare allosteric effect. Our results show that N23, N44, and N47 bind overlapping regions, as did NTD24 and NTD32 (**Figure 3B**). N42 bound a unique epitope.

**Figure 3:**
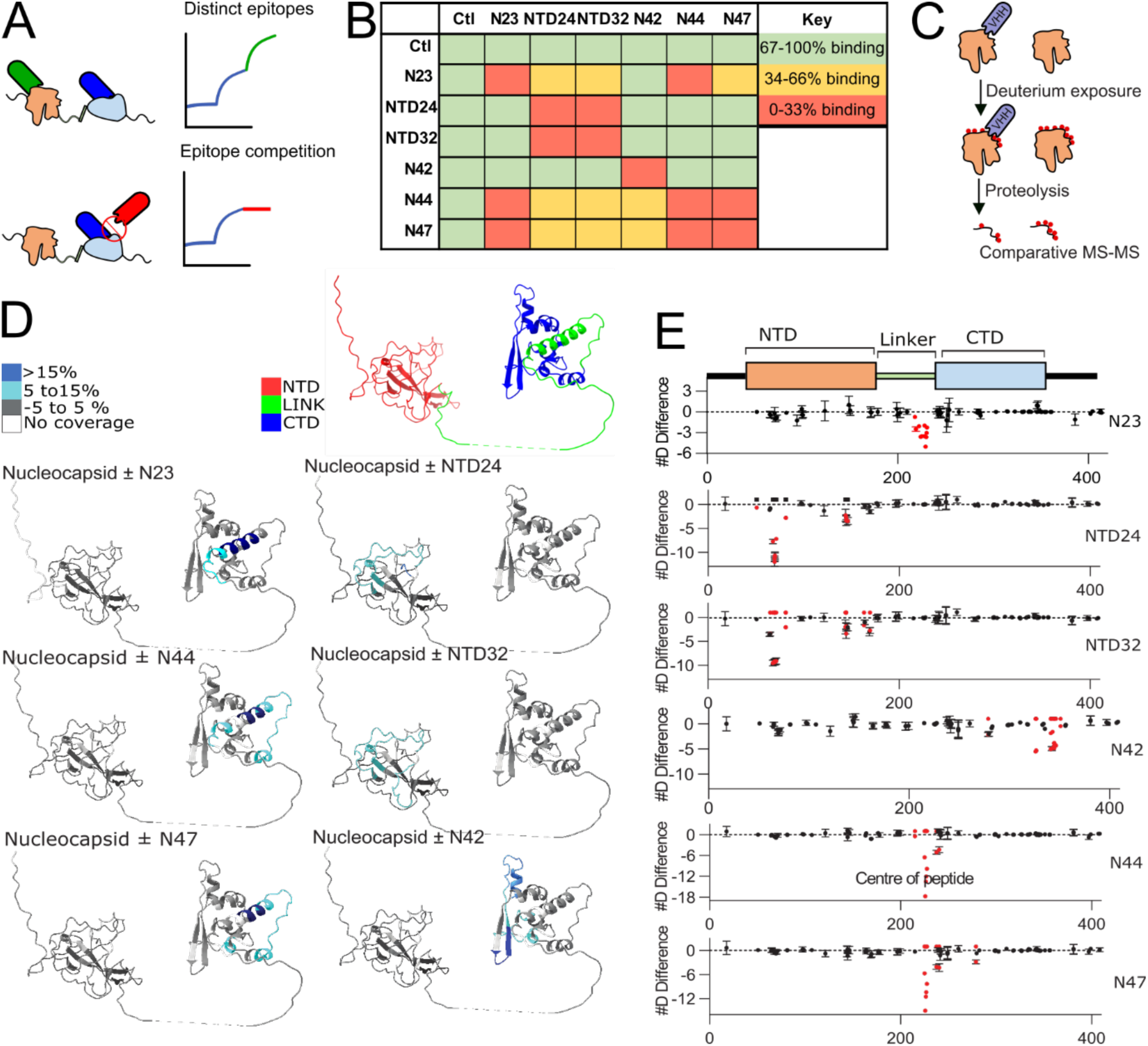
SARS-CoV-2 N VHHs specifically bind epitopes within the NTD, CTD, and LINK domains. A) Schematic for BLI competition assay. N is exposed to two VHHs in sequence. VHHs that do not compete for an epitope will result in two association curves (top panel) while those that do will yield one (bottom panel). B) Heatmap of competition potential between each VHH pair. The percent binding of an association used to bin VHH interactions into non-competitive (green), partially competitive (yellow) and competitive (red). C) Schematic for hydrogen-deuterium exchange (HDX-MS) to determine target domains for each VHH. Purified N in presence and absence of each VHH exposed to deuterium, fragmented, and ran through mass-spectrometry. D) Coverage map of areas showing significant differences in deuterium incorporation when comparing the apo SARS-CoV-2 Nucleocapsid to Nucleocapsid bound to each VHH. Changes are color coded according to the legend and have been mapped onto an AlphaFold3 model of the SARS-CoV-2 Nucleocapsid. E) Sum of the #D difference in deuterium incorporation upon addition of each VHH over the entire exchange time course. Each point represents a single peptide (n=3 error shown as SD) with significant changes in HDX (defined as >5%, >0.5 Da difference and a two-tailed t-test p < 0.01) highlighted in red. Nucleocapsid domain architecture is annotated above top graph.

To obtain higher resolution binding information, we next carried out HDX-MS of our best-binding VHHs on full-length N (**Figure 3C**). HDX-MS measures the rate of exchange of amide hydrogens for deuterium when incubated in a deuterated buffer. This rate of exchange for a given amide will be primarily dependent on the stability of protein secondary structure and solvent accessibility making it a well-suited technique to the mapping of protein-protein interactions and defining the intrinsic dynamics of protein structure^30^, with it being used extensively to map nanobody epitopes^31^. Full-length N was exposed to deuterium in the presence and absence of each VHH. The deuterium-labeled N was then digested into peptides and analyzed by mass spectrometry. Regions with decreased deuterium exchange in the presence of VHH compared to the apo nucleocapsid represent potential binding epitope regions. Our findings are summarized in (**Figure 3D-E**), with the full HDX-MS exchange information provided in the source data. N23, N44, and N47 had a similar region of decreased exchange in the presence of VHH; an alpha helix between amino acid residues 200 and 240, squarely in the LINK region. While N44 and N47 showed almost exactly the same differences in deuterium exchange, there was a different profile for N23, likely suggesting a similar but slightly distinct epitope. Exchange profiles of VHH NTD24 and NTD32 were mostly identical, identifying a region between residues 50 and 100 and 130-160 in the NTD, indicative of a shared epitope. HDX-MS results of VHH N42 show a potential epitope in the CTD, on a beta sheet between residues 325 and 375. Overall, the HDX-MS outcomes agree with the BLI competition assay, suggesting that we have obtained VHHs binding to each of the major N domains: NTD, LINK, and CTD.

To gain structural insights into VHH-N interactions, we used HDX-MS data to guide *in silico* modeling with AlphaFold3^32^. We applied two stringent criteria for evaluation of modeling results: (1) the predicted binding interface had to be supported by both HDX-MS and BLI-based VHH competition data, and (2) the complementarity-determining region 3 (CDR3) of each VHH had to serve as the primary interface with the N protein. We modeled N23, NTD24, NTD32, N42, N44, and N47 with the corresponding N domain identified by HDX-MS as the dominant binding site (**Figure 4A–F**). For NTD-specific VHHs NTD24 and NTD32, modeling was performed with 1 VHH and 1 copy of the NTD domain. For CTD-specific VHH N42, modeling was performed with 1 VHH and 2 copies of CTD, due to the well-established formation of stable CTD dimers^13,14,33,34^. For LINK-specific VHHs N23, N44, and N47, modeling was performed with 1 VHH and 2 copies of LINK+CTD because of the reported interactions between the LRH, to which these VHHs bind, and the CTD dimer^33^. While these *in silico* models should be interpreted with caution, AlphaFold3 consistently predicted (5 out of 5 times for all six VHHs) interactions that aligned well with HDX-MS data.

**Figure 4:**
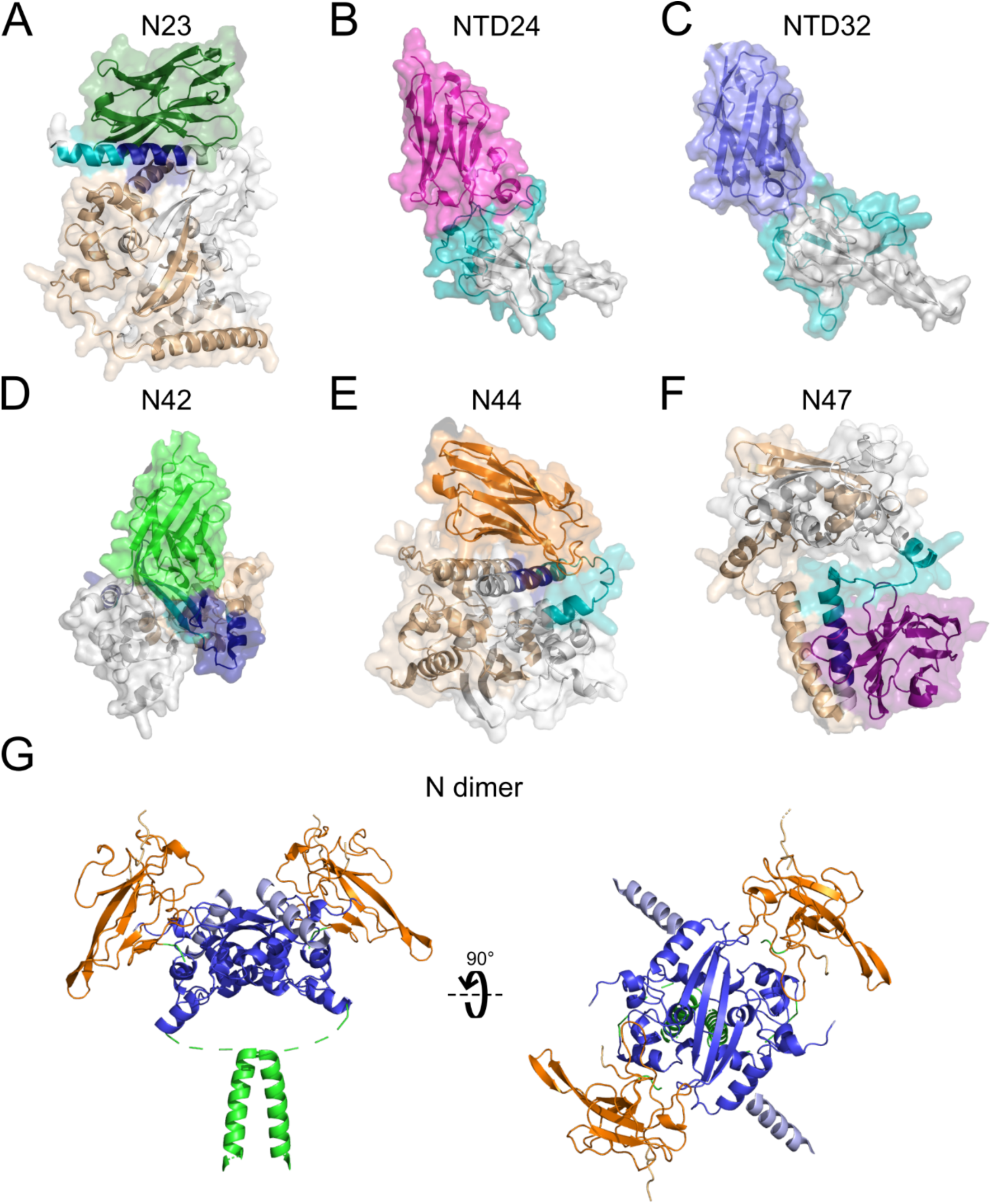
Modeling of nanobody binding. In silico modeling using AlphaFold3 of nanobody binding to N. A) N23 (green) binding to dimer of LINK+CTD. B) NTD24 (magenta) binding NTD. C) NTD32 (blue) binding NTD. D) N42 (green) binding dimer of CTD. E) N44 (orange) binding dimer of LINK+CTD. F) N47 (purple) binding dimer of LINK+CTD. N fragments indicate HDX difference data, Blue >15%, teal = 5-15%, white <5%. Secondary dimer component shown in tan. G) Modeling of the N dimer in complex with 2 copies of Sel15 RNA. N-arm in light orange, NTD in orange, LINK in green, CTD in blue, C-arm in light blue, RNA in purple. Low confidence regions (pLDDT<50) have been omitted from the model for clarity.

In order to see how the binding epitopes of these VHHs may impact N function, we next modeled N as a dimer and tetramer in 1:1 complex with an RNA 15mer (**Figure 4G-H and Supplemental Figure 3**). Previous studies have solved separate structures for the CTD and the NTD, but so far none have been able to obtain a structure of the full N protein due to the large disordered regions and high flexibility. Several studies have also used NMR to understand how RNA binds to the NTD^23,35^, finding that phosphorylation of the N-terminal portion of LINK has important effects on RNA binding and LLPS. NMR studies have also sought to understand the LINK region, finding that the LRH tends to form dimers (stabilized by CTD dimerization), which can then self-associate into tetramers under certain conditions^33^. Based on the modeling of our NTD-specific VHHs, NTD24 and NTD32, we would not expect abrogated RNA binding or LLPS. However, as the LRH directly interacts during dimerization and tetramerization, we would hypothesize that the LINK-specific VHHs N23, N44, and N47 may disrupt LLPS. Several studies have investigated which features of the N protein are important for ribonucleosomes formation, and while there are no high-resolution structures yet, it is clear that several regions including the LRH, and the CTD are indispensable for the formation of large oligomers^24,33,36,37^. Based on our modeling, we would not expect the CTD-specific VHH to disrupt dimerization, but it is not clear how this may impact formation of larger oligomers.

### N-specific VHHs inhibit viral replication through distinct mechanisms

We next wanted to empirically measure how each of our VHHs affect N function and viral propagation. To assess VHH impacts on viral replication, we created A549-ACE2 cells lines that stably expressed each VHH or irrelevant VHH which bound HIV matrix protein, with doxycycline (DOX) inducible control. We then infected each line with WA1, harvested progeny virus after 24 hours, and quantified progeny yields by focus forming assay (FFA) in VeroE6 cells (**Figure 5A-C**). All of the VHHs we tested (with the exception of N9) showed statistically significant inhibition of viral propagation, indicating functional inhibition of N by cytoplasmic VHHs. We confirmed that N9 was expressed at similar levels to the other VHHs in the A549 cells by quantifying the GFP intensity per cell (**Figure 5B**). The lowest expressing VHHs inside cells, NTD24 and NTD32, were still able to inhibit SARS-CoV-2 replication.

**Figure 5:**
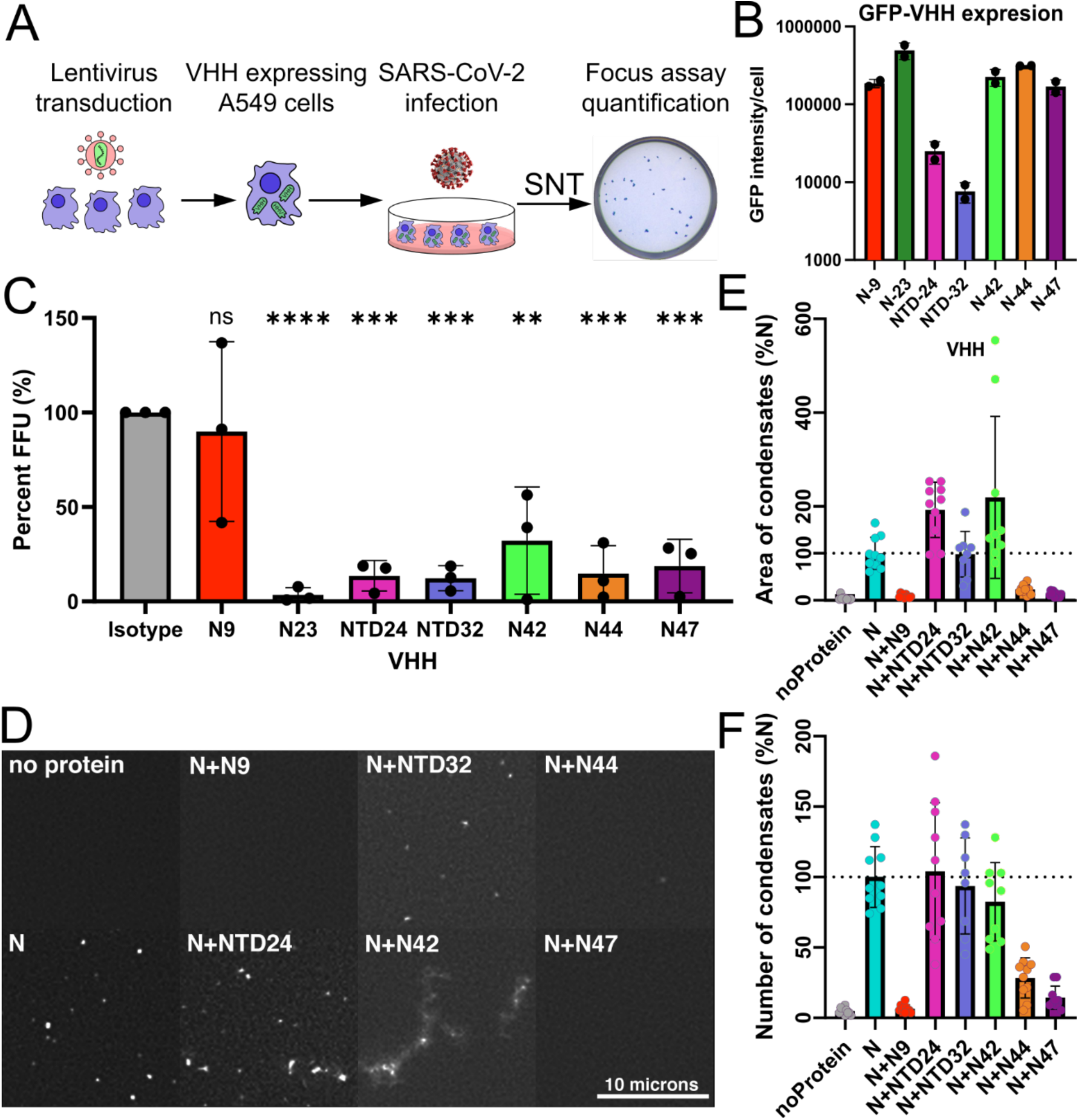
SARS-CoV-2 N-binding VHHs inhibit viral replication through distinct mechanisms, including LLPS. A) Schematic of SARS-CoV-2 infections. Lentivirus transduced VHH-expressing (cytoplasmic) A549 were infected with SARS-CoV-2 at an MOI of 1.0. Supernatant was taken at 24 hours post infection (HPI) and used in a focus forming assay (FFA) to quantify infectious virus. B) Quantitative microscopy of GFP signal to measure VHH expression level in GFP-VHH expressing stable cell lines, normalized by number of cells. C) FFU counts normalized to isotype control VHH. D) Fluorescent Sel15 RNA (3uM) alone, with N (5uM), and with the indicated VHH (15uM). Samples were applied to slides, incubated for 60 minutes, and imaged by fluorescence microscopy. E) Average numbers of RNA condensates from at least 8 independent images were determined as with standard deviations as indicated. F) Areas covered by RNA condensates from at least 8 independent images were determined as described in the methods section and were graphed after normalization to N protein-only sample, with standard deviations indicated.

To further probe the disruption of N function by VHHs, we next analyzed their ability to inhibit phase condensation with RNA, a critical step in viral genome packaging. We used a fluorescein-labeled RNA 15-mer and assessed condensate formation when mixed with recombinant N protein in the presence or absence of our domain-specific N VHHs. N9, N44, and N47 completely abrogated condensate formation, while N42 produced an altered phenotype that may reflect enhanced or aberrant condensate assembly (**Figure 5D**). VHHs NTD24 and NTD32 did not alter condensates. Specifically, VHH N9, N44, N47 reduced the number and brightness of condensates, while VHH N42 increased condensate brightness but not numbers, whereas NTD24 and NTD32 did neither (**Figure 5E-F**). N9 inhibited LLPS but not propagation; NTD24, NTD32, and N42 inhibited propagation but not LLPS; N44 and N47 inhibited both propagation and LLPS. These findings reveal distinct and sometimes uncoupled roles of N domains in phase condensation and viral replication, highlighting the utility of domain-specific VHHs as precise tools to dissect multifunctional viral proteins.

## Discussion

In this study we have optimized a pipeline for anti-viral-protein VHH development, and applied that to SARS-CoV-2 replication through targeting N. We successfully isolated six low-affinity and seven high-affinity VHHs by generating multiple libraries from an N-immunized alpaca. The seven highest-binding candidates bind full-length N in ELISA, and BLI, as well as successfully functioning in cytoplasm of SARS-CoV-2 infected cells. We confirmed by BLI that our collection of VHHs bind at least 3 distinct epitopes within N. Using HDX-MS, we found that the epitopes for our VHHs were distributed between the NTD, LINK, and CTD. When cytosolically expressed, many of these VHHs inhibited viral replication, though there may be distinct inhibitory mechanisms for each paratope; however, this will require further investigation. The inhibition of LLPS appears to be a common, but not universal property of N-specific VHHs, and our functional testing indicated that blocking LLPS is sufficient but not necessary for inhibiting SARS-CoV-2 replication. This implies distinct inhibitory mechanisms which do not require disruption of LLPS.

We observed a variety of phenotypes, which we can then match to the observed binding and functional data. Only LINK-specific VHHs blocked LLPS in our testing, with the exception of N9, for which we could not determine a precise epitope. Based on its strong inhibition of LLPS, we would hypothesize that N9 does bind LINK, but the low affinity paired with strong LLPS inhibition and weak to no inhibition of viral replication sets this VHH apart. The other interesting low affinity VHH, N23, stood out for its exceptional viral inhibition, and also its unique in silico modeling results in which it prevented dimerization of the LINK domains. We hope that future studies exploring the similarities and differences between N9-like and N23-like VHHs will help identify properties that distinctly inhibit viral replication versus LLPS among LINK-specific-VHHs.

Looking at the other domain of N, we observed that CTD-specific N42 was less effective at inhibiting replication, but it generated a unique result in the LLPS assay with larger and more dispersed clumping. Finally, NTD24 and NTD32 bind similar regions of the NTD and reduced viral progeny without affecting LLPS. Together, our findings provide further evidence that LINK plays an essential role in LLPS, but also that impairing LLPS *in vitro* does not guarantee inhibition of viral replication.

Our results also showed that NTD-binding was insufficient to block LLPS, while still blocking replication. However, a previous study on nanobodies against SARS-CoV-2 N showed that some NTD-specific VHHs are able to partially block LLPS, as well as interfering with RNA binding^14^. Those VHHs bound to a similar site as our VHHs NTD24 and NTD32, which could indicate that our results differed due experimental setup rather than epitope. In either case, the substantial difference between LINK-specific and NTD-specific VHHs in our LLPS assay leads us to believe that there is likely an alternative inhibitory mechanism to our LINK-specific VHHs. Higher resolution structural information about ribonucleosome assembly and packing may help develop testable hypotheses about such mechanisms.

Other studies have reported the use of nanobodies against N from SARS-CoV-2 as diagnostic detection reagents^39–41^, and we believe that our VHHs will also be suitable for this purpose. The N protein is more highly conserved than S, making it a generally better candidate for diagnostic development, and N is also produced in much greater quantities during infection^42^. The observed reactivity of SARS-CoV-1 antibodies against SARS-CoV-2 indicates that a future sarbecovirus is likely to be detectable using SARS-CoV-2-specific antibodies and nanobodies, and a greater breadth of available reagents will maximize the chance of success on this front while also providing critical resources for continued basic research into this growing family of viruses^43^.

Overall, we have demonstrated that VHHs are capable of impeding SARS-CoV-2 infection, that the NTD, CTD, and LINK are all effective targets for an inhibitory antibody fragment, and that binding the LINK alone is sufficient to impede RNA-N LLPS. Further structural work will help confirm the involvement of individual residues and elucidate functional mechanisms. While the use of Alphafold3 for structural prediction is a limitation of the study, we have validated it empirically against our HDX-MS data and we believe there is value in using this approach for generating hypotheses. We hope that our clear findings on the susceptibility of the LINK domain to disruption by nanobodies will encourage further investigation of LINK targeting therapeutics for SARS-CoV-2, as well as anti-viral nanobody projects against similar antigens from other viruses.

## Materials & Methods

### N protein purification and biotinylation

N protein was expressed in PlysS DH5-α *E. coli* and was isolated cytosolically by ammonium sulfate precipitation. *E. coli* containing a codon-optimized his-tagged N pET-24a vector were grown in Luria Broth supplemented with10 mM MgSO_4_ and 0.02% maltose and were induced with 1mM IPTG and grown at 37°C for 4 hours, then centrifuged at 4000 rcf at 4°C for 10 minutes and frozen at -80°C. Pellets were solubilized in wash buffer (10mM Tris 500mM NaCl pH 7.5) supplemented with 20 µg/mL DNAse and 10 µg/mL RNAse to reduce N-nucleotide interactions, then cells were lysed by French press. Lysate was clarified by centrifugation at 20,000 rcf at 4°C for 20 minutes. Supernatants were treated with 0.3% PEI to remove nucleotides, then clarified again. The nucleotide-free supernatant was precipitated with 60% saturation volume ammonium sulfate, then pelleted at same conditions. Protein pellets were resuspended in wash buffer, then purified by Ni-NTA chromatography, washed with wash buffer with 20mM imidazole, and eluted with wash buffer containing 250 mM imidazole. Purified protein was then buffer exchanged into storage buffer (500 mM NaCl, 50 mM NaPO_4_, 5 mM BME and 10% glycerol, pH 8), then frozen at -80°C. For use in BLI, purified N was biotinylated using the ChromaLINK biotin protein labeling kit (Vector) according to the manufacturer’s instructions with 5x molar equivalents of labeling reagent to achieve ∼2 biotin/protein.

### VHH gene library construction

To isolate VHH sequences against SARS-CoV-2 N, first immunized an alpaca with recombinant full-length N protein isolated as described above. Protein purity was screened by SDS-PAGE. Alpacas were immunized at Capralogics Inc. (Hardwick, MA). Immunization consisted of 1 mg purified full-length N in sterile PBS with Imject alum adjuvant (Thermo Fisher scientific) administered intramuscularly in alternating shoulders. One round consisted of four immunizations spaced three weeks apart and blood was harvested five days after the fourth injection. Two rounds of immunizations were carried out approximately 1 year apart. Upon receipt, PBMCs were isolated with Ficol paque (Cytvia life sciences) as described previously^44^ and subjected to RNA extraction with Qiagen RNeasy mini kit (Qiagen). PBMC RNA, containing VHH genes, was converted into cDNA using Superscript III reverse transcriptase (Invitrogen). VHH genes were amplified with custom primers, as published previously^45^, that were specific to short-hinge and long hinge VHH genes appended with Not1 and Asc1 restriction sites. The amplified gene mixture was cloned into a phage-mid plasmid derived from pCANTA5BE with these restriction sites. The 200ng of pooled library DNA in water was then then transformed via electroporation into 50 µL of bacteriophage-sensitive TG1 *E. coli* and recovered in SOB media at 37°C to produce of a VHH displaying bacteriophage library, which was plated. Vector-only and insert-only controls were included to confirm background was lower than 0.5% of the total library count. Plates containing serial dilutions were used to estimate total bacterial population, and therefore library diversity; 1×10^7^ and 3×10^7^ cfu for the first and second immunization course libraries, respectively. A random sample of 96 VHH clones were sent for sanger sequencing

### VHH panning

M13-derived helper phage was produced through standard protocols described previously^46^. VHH libraries in TG1 cells were transduced with helper phage to produce bacteriophage displaying individual VHHs. Phage were isolated through precipitation using 20% w/v polyethylene glycol 8000, then were resuspended in PBS. Phage libraries were then panned against full length N or N NTD domain at 20 µg/ml alongside BSA controls. Bacteriophage were bound to the antigen, washed, then eluted with 200 mM glycine pH 2.2, neutralized with 1M Tris pH 9.1, and transduced into ER2738 bacteriophage competent *E. coli* which were plated on antibiotic agar and incubated for 18 hours at 37°C. Panning success was determined by enrichment 10-fold greater total bacterial colonies above control panning background. Panning colonies were then pooled, and the protocol was repeated for second round panning using 2 µg/ml antigen coating. A selection of colonies from second round panning were picked and grown up in 96 deep-well plates for screening. Screening of VHH candidates 96 well ELISAs coated with 1 µg/ml purified N were used to determine N-binding affinities of panning hits. ELISAs were performed with bacterial supernatant containing secreted VHHs as primary antibody, followed by a secondary anti-nanobody biotin antibody (Jackson #128-065-232) and streptavidin-HRP (Thermo #N100). 3,3’5,5’-tetramethylbenzidine (TMB) (ThermoFisher Scientific) was used as peroxidase substrate, 50 mL added for 10 minutes at room temperature (RT) then 50 mL of H_2_SO_4_ was added as a stopping solution. Plate absorbance at 405 nm was measured using a CLARIOstar Plus plate fluorimeter (BMG Labtech). VHH candidates with binding greater than 2-fold above average background were picked and sent for Sanger sequencing (Genewiz) to identify VHH CDR3 regions. Sequences with homology were considered conserved families.

### VHH purifications

Clones that appeared multiple times in sequencing were cloned into a periplasmic expression vector (pHEN) by Gibson assembly standard protocol (NEW England biolabs). Bacteria were grown up in terrific broth (2% tryptone, 1% yeast extract, 90mM phosphate), induced with IPTG at 30°C overnight. VHHs were purified by periplasmic expression and osmotic shock was used to isolate the periplasmic fraction which contain 6xHis tagged protein, VHHs were purified with Ni-NTA chromatography. Purified VHH was then buffer exchanged and concentrated with 3 KDa cutoff centrifuge filters (Millipore), then sterilized by 22 nm centrifugal sterile filter (Millipore Sigma) prior to use in experiments.

### Enzyme-linked immunosorbent assay (ELISA)

MaxiSorp ELISA plates (Invitrogen), were coated with purified recombinant full-length SARS-CoV-2 N 2 mg/mL in PBS, Coating was carried out overnight at 4°C. Plates were blocked in blocking buffer (2% BSA, 0.1% tween-20 in PBS) for 30 minutes at RT. Dilutions of N VHHs in blocking buffer were incubated for 1 hour at RT. Plates were washed with PBST (0.1% tween-20 in PBS) 4 times between each antibody addition. Anti-VHH-biotinylated antibody and streptavidin-HRP secondary antibodies were used at 1:10000 concentration in blocking buffer and were incubated 1 hour at RT. After the final wash, plates were incubated for 10 minutes with 50 mL of TMB (ThermoFisher) at RT, before adding 50 mL of stopping solution (2N H_2_SO_4_). Plate absorbances at 405nm were measured using a CLARIOstar Plus plate fluorimeter (BMG Labtech).

### Biolayer interferometry (BLI)

Streptavidin biosensors (Sartorius) were soaked in PBS for at least 30 minutes prior to starting experiments. Biosensors were prepared with the following steps: equilibration in kinetics buffer (10 mM HEPES, 150 mM NaCl, 3mM EDTA, 0.005% Tween-20, 0.1% BSA, pH 7.5) for 300 seconds, loading of biotinylated N protein (10 µg/mL) in kinetics buffer for 200 seconds, and blocking in 1 mM D-Biotin in kinetics buffer for 50 seconds. Binding was measured for seven 3-fold serial dilutions of each monoclonal antibody using the following cycle sequence: baseline for 300 seconds in kinetics buffer, association for 300 seconds with antibody diluted in kinetics buffer, dissociation for 750 seconds in plain kinetics buffer, and regeneration by 3 cycles of 20 seconds in 10 mM glycine pH 1.7, then 20 seconds in kinetics buffer. All antibodies were run against an isotype control antibody at the same concentration.

For competition experiments, biosensors were loaded with full-length N similarly to binding experiments, then bound with 400 nM of each antibody for 300 seconds before transferring to a competing antibody diluted in kinetics buffer for 300 seconds. Data analysis was performed using the ForteBio data analysis HT 10.0 software. Curves were reference subtracted using the isotype control and each cycle was aligned according to its baseline step. *K_D_*s were calculated using a 1:1 binding model, and the kinetic parameters (*K_D_*, *k_ON_*, *k_OFF_*) were averaged from concentrations and replicates, excluding dilutions with an R^2^ less than 0.9 or an Rmax more than double the average of other concentrations.

### Hydrogen-deuterium exchange mass spectrometry (HDX-MS)

HDX-MS reactions comparing SARS-CoV-2 N unbound to the NTD24, NTD32, N42, N44, and N47 bound states were carried out in 30 µL reaction volumes containing 15 pmol of protein. The exchange reactions were initiated by the addition of 26 µL of D_2_O buffer (20 mM HEPES pH 7.5, 100 mM NaCl, 94.34% D_2_O [v/v]) to 4 µL of protein (final D_2_O concentration of 81.7% [v/v], 500 nM Nucleocapsid, 1 µM nanobody). HDX-MS reactions comparing SARS-CoV-2 N unbound to the N23-bound state were carried out in 50 µL reaction volumes containing 25 pmol of protein. The exchange reactions were initiated by the addition of 43.44 µL of D_2_O buffer to 6.66 µL of protein (final D_2_O concentration of 81.7% [v/v], 500 nM Nucleocapsid, 1 µM nanobody). Reactions proceeded for 3s, 30s, 300s, or 3000s before being quenched with ice cold acidic quench buffer, resulting in a final concentration of 0.6M guanidine HCl and 0.9% formic acid post quench. All condition and timepoints were created and run in independent triplicate. Samples were flash frozen immediately after quenching and stored at -80°C until injected onto the ultra-performance liquid chromatography (UPLC) system for proteolytic cleavage, peptide separation, and injection onto a QTOF for mass analysis, described below.

### Protein digestion and MS/MS data collection

Protein samples were rapidly thawed and injected onto an integrated fluidics system containing an HDX-3 PAL liquid handling robot and climate-controlled (2°C) chromatography system (LEAP Technologies), a Dionex Ultimate 3000 UHPLC system, as well as an Impact HD QTOF Mass spectrometer (Bruker). The full details of the automated LC system as previously described^47^. The unbound, NTD24, NTD32, N42, N44, and N47 bound samples were run over one immobilized pepsin column (Trajan; ProDx protease column, 2.1 mm x 30 mm PDX.PP01-F32) at 200 µL/min for 3 minutes at 8°C. The second unbound and N23-bound samples were run over two immobilized pepsin columns (Waters; Enzymate Protein Pepsin Column, 300Å, 5µm, 2.1 mm X 30 mm) at 200 µL/min for 3 minutes at 2°C. The resulting peptides were collected and desalted on a C18 trap column (Acquity UPLC BEH C18 1.7mm column (2.1 x 5 mm); Waters 186003975). The trap was subsequently eluted in line with an ACQUITY 1.7 μm particle, 100 × 1 mm2 C18 UPLC column (Waters), using a gradient of 3-35% B (Buffer A 0.1% formic acid; Buffer B 100% acetonitrile) over 11 minutes immediately followed by a gradient of 35-80% over 5 minutes. Mass spectrometry experiments acquired over a mass range from 150 to 2200 m/z using an electrospray ionization source operated at a temperature of 200°C and a spray voltage of 4.5 kV.

### Peptide identification

Peptides were identified from a non-deuterated SARS CoV2 Nucleocapsid apo sample using data-dependent acquisition following tandem MS/MS experiments (0.5 s precursor scan from 150-2000 m/z; twelve 0.25 s fragment scans from 150-2000 m/z). MS/MS datasets were analysed using PEAKS7 (PEAKS), and peptide identification was carried out by using a false discovery-based approach, with a threshold set to 0.1% using a database of purified proteins and known contaminants^48^. The search parameters were set with a precursor tolerance of 20 ppm, fragment mass error 0.02 Da, charge states from 1-8, leading to a selection criterion of peptides that had a -10logP score of 32.6.

### Mass analysis of peptide centroids and measurement of deuterium incorporation

HD-Examiner Software (Sierra Analytics) was used to automatically calculate the level of deuterium incorporation into each peptide. All peptides were manually inspected for correct charge state, correct retention time, appropriate selection of isotopic distribution, etc. Deuteration levels were calculated using the centroid of the experimental isotope clusters. Results are presented as relative levels of deuterium incorporation and the only control for back exchange was the level of deuterium present in the buffer (81.7%). Differences in exchange in a peptide were considered significant if they met all three of the following criteria: ≥5% change in exchange, ≥0.5 Da difference in exchange, and a p value <0.01 using a two tailed student t-test. Samples were only compared within a single experiment and were never compared to experiments completed at a different time with a different final D_2_O level. The data analysis statistics for all HDX experiments are in Supplemental Source Data according to published the guidelines^49^. The mass spectrometry proteomics data have been deposited to the ProteomeXchange Consortium via the PRIDE partner repository with dataset identifier PXD043179^50^.

### Cell culture

Low-passage HEK-293T, HEK-293T-ACE2, and Vero E6 cells were cultured in D10, which consisted of DMEM supplemented with 10% fetal bovine serum (FBS), 1% Penn-Strep, 1% non-essential amino acids (NEAA). 37°C. A549-ACE2 were grown in F12-K media with 10% FBS. Selection F12-K media was modified with 1% Doxycycline, 700 mg/ml G418 was used for stable cell lines. Cells were cultured in T75 dishes, passaged with Trypsin at 95% confluency to avoid overcrowding. Cell lines were frozen down in freezing media consisting of 5% DMSO, 95% FBS.

### SARS-CoV-2 virus propagation

Clinical isolation of SARS-CoV-2 WA1/2020 (NR-52281) was obtained via BEI resources and propagated in our BSL3 laboratory. To propagate, a 70% confluent T25 flask of Vero E6 cells was infected at a MOI of 0.01 in diluted in 1 mL Opti-MEM for 1 hour at 37°C with occasional rocking. 4mL of D10 was then added and the flask was incubated for 72 hours at 37°C. Following incubation, flasks were checked for cytopathic effect (CPE), after which supernatant was collected and spun at 3000 rcf for 5 minutes to remove cellular debris, then aliquoted for storage at −80°C. Propagated stocks were titrated with 8 × 10-fold dilutions in a focus forming assay as described below.

### Focus forming assay (FFA) of stable cell lines

A549-ACE2 lines for each VHH were seeded at 10,000 cells/well in presence of 1µg/ml doxycycline to stimulate VHH expression. They were then infected with an MOI of 1.0 of WA1/2020 SARS-CoV-2. Cells supernatants were then harvested 24 HPI and cells were fixed in 4% formaldehyde for use in florescent microscopy. For focus forming assay, Vero E6 cells were plated at 20,000 cells/well in 96-well plates and incubated overnight. They were then infected with neat infectious supernatant or 10-fold dilutions in Optimem, then incubated for 1 hour at 37°C while rocking. Afterwards, 150 mL of overlay media (Opti-MEM, 2% FBS, 2% Methylcellulose) was then added to each well and incubated for 24 hours at 37C. Plates were fixed using 4% PFA for 1 hour at RT. Plates were then blocked for 30 minutes with permeabilization buffer (PBS, 0.1% BSA and 0.1% saponin). RBD-immunized-alpaca^51^ sera was used as a primary antibody at 1:5,000 dilution in permeabilization buffer, and anti-Llama-HRP secondary was used at 1:20,000 dilution in permeabilization buffer. Plates were developed in 30 µL TrueBlue (SeraCare) substrate and imaged with an Immunospot analyzer (CTL). Foci were counted with Viridot version1.0 in R version 4.1.0.

### SARS-CoV-2 immunofluorescence

For testing of VHH candidate binding of infected cells, 96-well TC plates were seeded to 50% confluency with VeroE6 cells. Plates were then inoculated at a MOI of 0.1 of SARS-CoV-2 in 50 μL of Opti-MEM for 1 hour at 37°C with occasional rocking. An additional 50 μL of fresh media was then added and incubated for 24 hours at 37°C. Plates were fixed by submerging in 4% paraformaldehyde (PFA) in PBS for 1 hour, then brought into BSL-1 for immunofluorescence staining. Permeabilization was performed with 2% bovine serum albumin (BSA), 0.1% Triton-X 100 in PBS for 1 hour at RT. Cells were stained with each N VHH at 2 μg/mL, and mouse anti-dsRNA (Millipore sigma # MABE1134) 1:1000. Anti-llama IgM AF488-conjugated secondary antibodies and anti-mouse-AF555-conjugated antibody were added at 1:500 dilution for 1 hour at RT (Invitrogen). Confocal imaging was performed with a Zeiss LSM 980 using a 63x Plan-Achromatic 1.4 NA oil immersion objective. Images were processed with Zeiss Zen Blue software. Maximum intensity *z*-projections were prepared in ImageJ. All antibody stain images were pseudo-colored for visual consistency.

### Phase condensation assay

Fluorescein isothiocyanate-labeled Sel15 RNA (FITC-Sel15; sequence: FITC-GGAAU UAAUA GUAGC) was added to one of three conditions: buffer alone, recombinant full-length N protein (expressed in *E. coli*), or N protein preincubated with anti-N VHHs. VHHs and N protein were preincubated for 10 minutes at 25°C prior to RNA addition. Final sample volumes were 10 µL, with final concentrations of 3 µM FITC-Sel15 (45 µM total nucleotide), 5 µM N protein, and 15 µM VHHs, in a buffer containing 50 mM sodium phosphate (pH 6.0), 162.5 mM NaCl, 5 mM Tris, and 1.25 mM β-mercaptoethanol (BME). Samples were applied to microscope slides and covered with 18 mm diameter, 0.15 mm thick coverslips (Fisher 18CIR-1), sealed with clear nail polish, and incubated for 60 minutes at 25°C before imaging. Imaging was performed on a Zeiss AxioObserver fluorescence microscope using a 63x Plan-Apochromat objective and Zeiss filter set 10 (excitation: 450–490 nm; beamsplitter: 510 nm; emission: 515–565 nm). Images were acquired as TIFF files using Zeiss Axiovision software, with an exposure time of 100 ms and a fixed gain of 100.

### Fluorescent microscopy analysis and quantitation

For fluorescent analysis of SARS-CoV-2 infection, total surface area and intensity of blue and red signal were quantified using Keyence software. Three technical replicates were performed for each concentration in each experiment. The average of the triplicates red signal (anti DS-RNA) was normalized to average blue signal (DAPI) for each concentration. Resulting values were then subtracted with the red signal from uninfected control, then normalized to the infected control signal and plotted. Pairwise ANOVA was used for two experiments to determine significance.

For quantitation of infection of stable cell lines, fixed, infected A549-ACE2 cell lines, as described above, were stained with DAPI and mouse anti-dsRNA (Millipore sigma # MABE1134) 1:1000 and anti-mouse-AF555-conjugated secondary antibody. Cells were imaged with BZ-X700 all-in-one fluorescent microscope (Keyence). Estimated area of DAPI and AF555 pixels were calculated with built in BZ-X software as described below (Keyence).

For analysis of RNA-protein condensates, 3600 square micron areas from at least 8 independent images for each sample were thresholded to image brightness means plus four standard deviations using the Fiji (Image J) software Image/Adjust/Threshold command. RNA condensates were picked using the Fiji Analyze/Analyze Particles command, with size parameters of 0 to infinity pixels, and circularity parameters of 0.0-1.0. Particle numbers were averaged for each sample and normalized to the RNA plus pet24-Sars-N-his only sample to obtain normalized number of condensate values. Total particle areas were averaged for each sample and normalized to obtain normalized area of condensate values. Note that raw RNA plus pET-24-SARS-CoV-2-N-His only values for 3600 square micron samples were 55.4 +/- 12.0 particles and 125.9 +/- 42.8 particle pixel areas. For statistical significance determinations, means and standard deviations were converted to Z values assuming normal distributions, and statistical significance values were calculated from Z values.

## Acknowledgements

BLI data were generated on an Octet Red 384, which is made available and supported by OHSU Proteomics Shared Resource facility and equipment grant number S10OD023413. We also thank the OHSU Flow Cytometry Shared Resource, and OHSU Advanced Light Microscopy Core for the use of their software, equipment, and expertise. This work was supported by the National Institutes of Health grant 1R01AI141549-01A1, the MJ Murdock Foundation (F.G.T.), and T32AI170496 (T.A.B.). J.E.B. is supported by the Natural Science and Engineering Research Council of Canada (NSERC) Discovery Grant (2020-0424). E.B. was supported by NIH grant R01AI152579.

## Supplemental figures and tables

**Supplemental table 1:** Summary of VHHs and Fc-dimers. Summary of VHH binding data including ELISA EC_50_, BLI *K_D_*, and HDX-MS epitope mapping. Bolded VHHs were taken forward for functional testing. “-” for no data, “ND” for not determined.

| VHH Monomers | Panning round | ELISA EC<sub>50</sub> (nM) | BLI K<sub>D</sub> (nM) | Domain specificity (HDX-MS) |
| --- | --- | --- | --- | --- |
| N5 | 1 | 2679 | ND | - |
| N6 | 1 | 1094 | ND | - |
| N8 | 1 | 538.5 | ND | - |
| **N9** | 2 | 150 | 1088 | ND |
| N16 | 1 | 57.1 | ND | - |
| **N23** | 1 | 273.9 | ND | LINK |
| **NTD24** | 2 | 0.07 | 0.063 | NTD |
| N31 | 1 | 2956 | ND | - |
| **NTD32** | 2 | 0.06 | 4.08 | NTD |
| **N42** | 2 | 0.22 | 2.61 | CTD |
| **N44** | 2 | 0.47 | 10.5 | LINK |
| **N47** | 2 | 0.4 | 8.40 | LINK |
| Fc-VHH dimers | - | ELISA EC<sub>50</sub> (nM) | BLI apparent K<sub>D</sub> (nM) | Domain specificity (HDX-MS) |
| FcN5 | 1 | 1.2 | 11.4 | - |
| FcN6 | 1 | 0.8 | 10.7 | - |
| FcN16 | 1 | 1.6 | 13.8 | - |
| FcN23 | 1 | 0.3 | 2.78 | LINK |
| FcN31 | 1 | 0.8 | 12.6 | - |

**Supplemental Figure 1:**
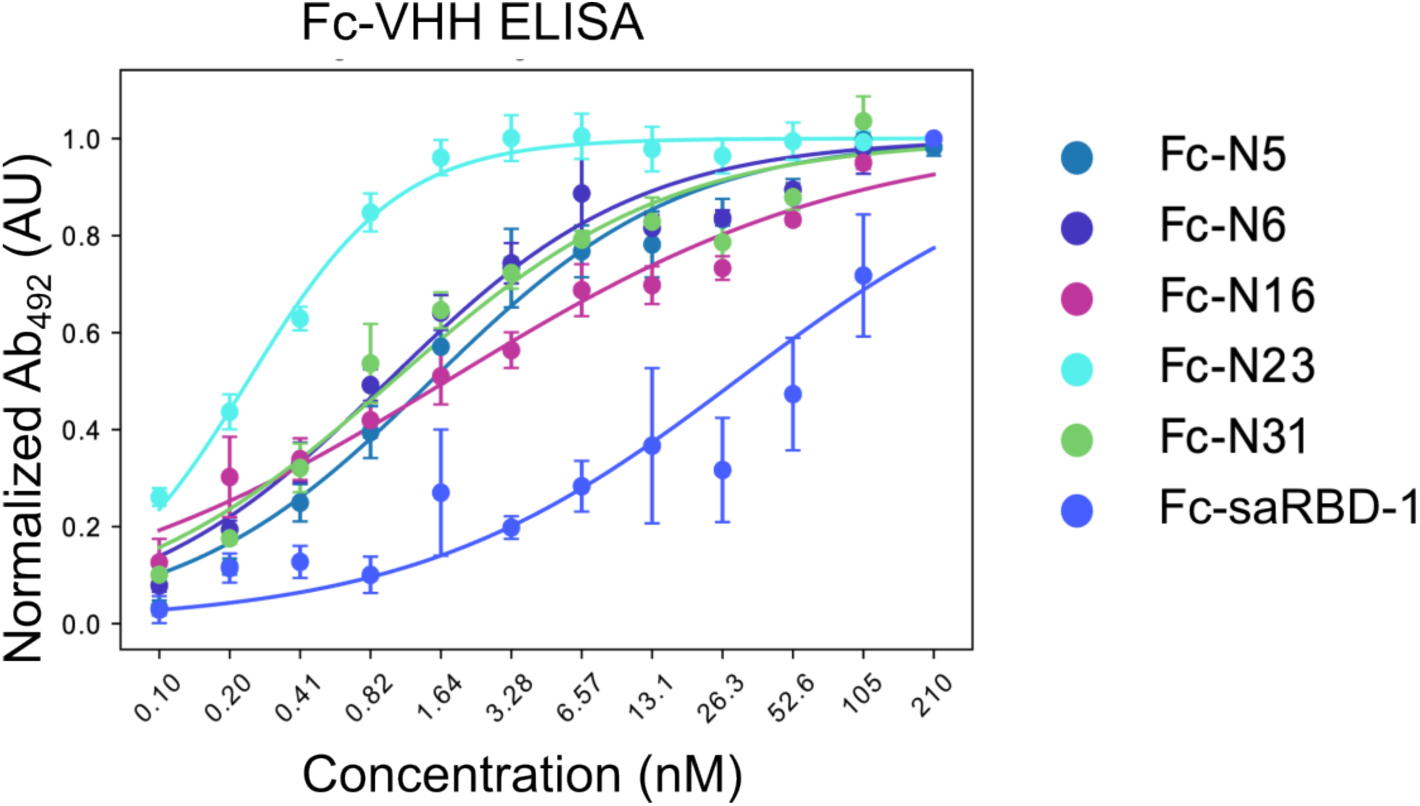
ELISA of Fc-dimers of N VHH candidates against full-length SARS-CoV-2 N protein. Bivalent Fc-VHH dimers were created by cloning VHH genes into an IgG1 backbone and expressed in 293-F cells. An anti-human IgG secondary antibody was used for detection. Spike-specific Fc-saRBD-1 built using the same IgG1 backbone was used as a control ELISA coated with spike receptor binding domain (RBD).

**Supplemental Figure 2:**
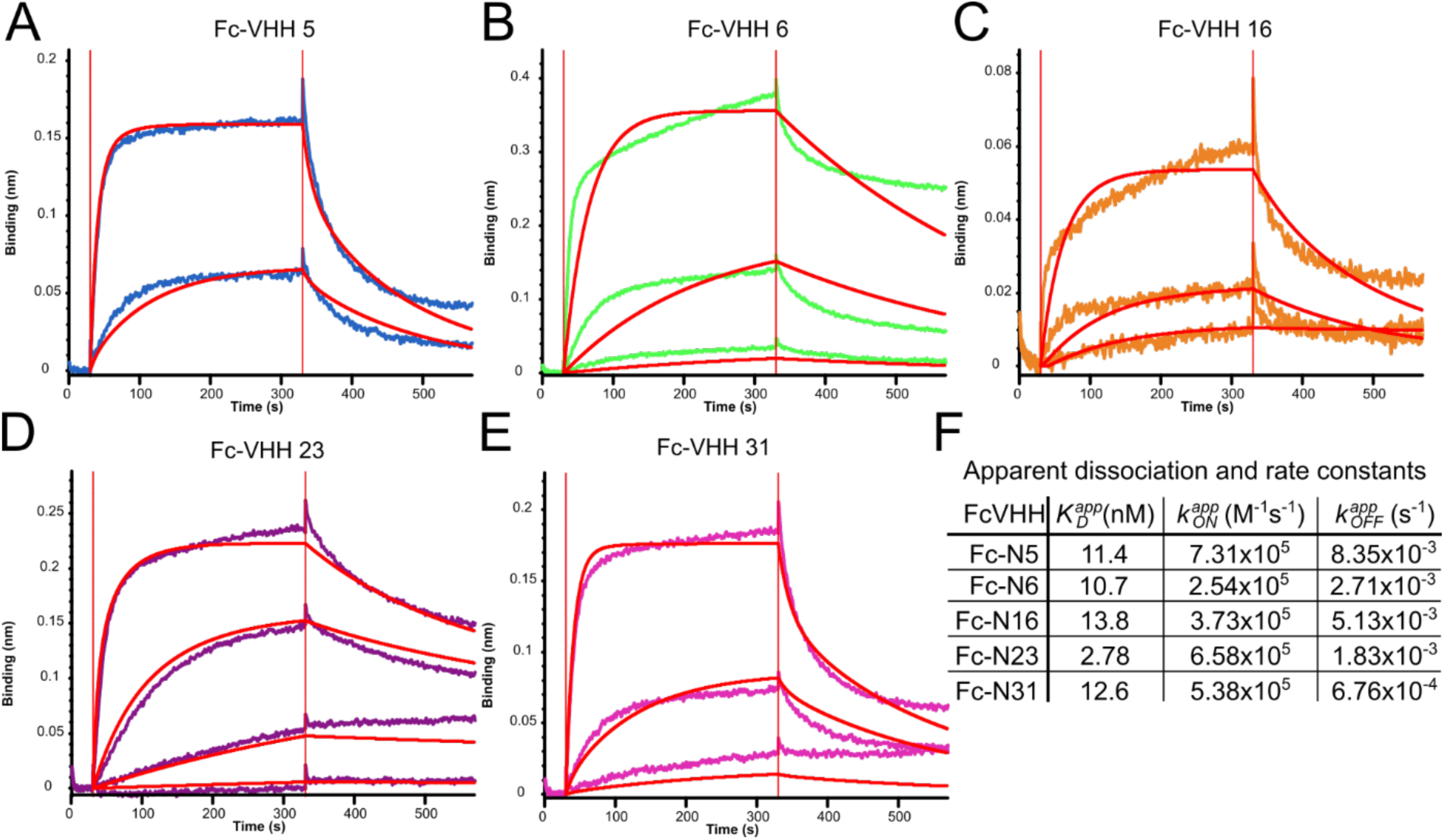
BLI of Fc-dimers of N VHH candidates against full-length SARS-CoV-2 N protein. N-coated biosensors were incubated with bivalent human IgG1 FC-nanobodies. A) Fc-VHH N5, B) Fc-VHH N6, C) Fc-VHH N16, D) Fc-VHH N23, E) Fc-VHH N31. F) Apparent *K_D_*, *k_ON_*, and *k_OFF_* for bivalent Fc-VHHs calculated using a 1:1 model to allow comparison with higher-affinity monomers.

**Supplemental Figure 3:**
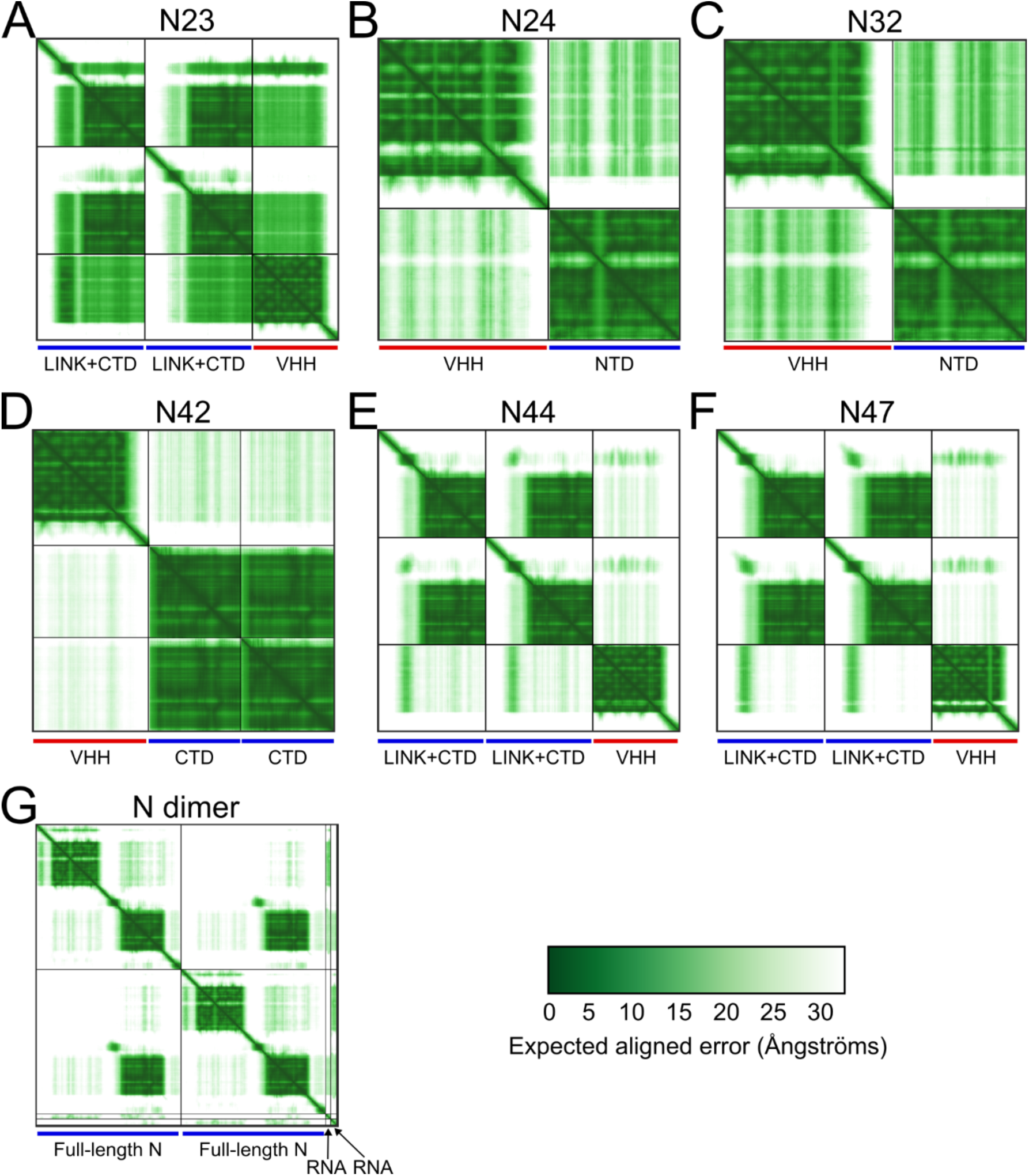
Predicted aligned error (PAE) plots for alphafold3 predictive models. PAE plots for the models shown in Figure 4. A) N23 was modeled on a dimer of LINK+CTD, B) NTD24 was modeled on a single NTD, C) NTD32 was modeled on a single NTD, D) N42 was modeled on a dimer of CTD, E) N44 was modeled on a dimer of LINK+CTD, F) N47 was modeled on a dimer of LINK+CTD, G) A dimer of full length N was modeled with 2 molecules of sel15 RNA.

